# Semantic harmonization of Alzheimer’s disease datasets using AD-Mapper

**DOI:** 10.1101/2023.10.26.564134

**Authors:** Philipp Wegner, Helena Balabin, Mehmet Can Ay, Sarah Bauermeister, Lewis Killin, John Gallacher, Martin Hofmann-Apitius, Yasamin Salimi, the Alzheimer’s Disease Neuroimaging Initiative, the Japanese Alzheimer’s Disease Neuroimaging Initiative, the Aging Brain: Vasculature, Ischemia, and Behavior Study, the Alzheimer’s Disease Repository Without Borders Investigators, the European Prevention of Alzheimer’s Disease (EPAD) Consortium

## Abstract

**INTRODUCTION:** Despite numerous past endeavors for the semantic harmonization of Alzheimer’s disease (AD) cohort studies, an automatic tool has yet to be developed. As cohort studies form the basis of data-driven analysis, harmonizing them is crucial for cross-cohort analysis. We aimed to accelerate this task by constructing an automatic harmonization tool.

**METHODS:** We created a common data model (CDM) through cross-mapping data from 20 cohorts, three CDMs, and ontology terms, which was then used to fine-tune a BioBERT model. Finally, we evaluated the model using three previously unseen cohorts and compared its performance to a string-matching baseline model.

**RESULTS:** Here, we present our AD-Mapper interface for automatic harmonization of AD cohort studies, which outperformed a string-matching baseline on previously unseen cohort studies. We showcase our CDM comprising 1218 unique variables.

**DISCUSSION:** AD-Mapper leverages semantic similarities in naming conventions across cohorts to improve mapping performance.

## 1. Background

In Alzheimer’s disease (AD) research, numerous cohort datasets serve as the foundation for data-driven investigations (e.g., based on machine learning (ML)). These datasets are often customized to address specific research questions and, therefore, focus on specific biomarkers and measurements that are essential for the research [1]. Such collected measurements are usually stored in different formats and using arbitrary naming systems. These inconsistent variable naming conventions and metadata across cohorts impede interoperability and make cross-cohort research time-consuming [2]. Despite the growing number of collected AD cohort datasets, harmonizing and utilizing multiple cohorts for disease investigation remains challenging due to these variable naming differences. As a result, the majority of research is practically limited to single cohorts. However, numerous reports indicate that conclusions drawn from AD data were constrained to the cohorts used and may not necessarily be generalizable [3, 4]. Therefore, single-cohort studies benefit from validation using independent datasets [5]. To address this and encourage cross-cohort investigations, it is vital to identify a common ground for AD data harmonization [6], ideally, using an automated tool.

Motivated by these matters, several attempts have been made to harmonize AD cohort studies by generating data catalogs, common data models (CDMs), and data stewardship tools (DSTs). Recently, Salimi *et al.* (2022) demonstrated the differences that exist concerning over 1000 collected measurements and the naming convention across 20 major AD cohort studies through manual curation. They harmonized the cohorts’ variables against normalized variable names in addition to ontology terms. Their endeavor established the foundation for implementing a harmonized AD landscape, aiding researchers in cohort data selection and ensuring data interoperability [2]. Similarly, Bauermeister *et al.* (2023) proposed the C-Surv data model, covering the harmonization of 124 variables across four distinct cohorts [7]. Alternatively, other attempts were made to establish a data catalog and patient/variable outcome. For instance, the ROADMAP data cube and the EMIF data catalog projects illustrate the data availability among multiple cohort studies [8, 9]. While both of these projects included many cohorts and modalities, the reported information was mainly gathered through the data owners and the corresponding metadata. Such a data catalog did not address the available variables on a granular level and the variable harmonization aspect across cohort studies. Another study by Wegner *et al.* (2022) established a semi-automatic DST using a string-matching technique for the harmonization of clinical datasets and applied it in the field of dementia [10]. However, despite previous efforts, there is currently no model or tool enabling fully automatic harmonization in the AD field. Consequently, data curation/harmonization has predominantly remained a manual task.

One promising direction to address the aforementioned issue is the automation of manual mappings using natural language processing (NLP). In recent years, approaches such as the Bidirectional Encoder Representations from Transformers (BERT) model [11] and its biomedical equivalent, BioBERT [12], have greatly improved the ability of language models to correctly represent a word or phrase using its surrounding context. As a result, these models offer more flexibility for mapping purposes compared to previous methods based on string matching. Further, to account for domain shifts, such pre-trained language models (PLMs) can be fine-tuned to a wide range of tasks using task-specific datasets [13]. Nonetheless, to the best of our knowledge, no approach has leveraged pre-trained biomedical language models for variable harmonization between AD cohort datasets yet.

Here, we implemented AD-Mapper, an automatic tool for the semantic harmonization of AD cohort datasets. We developed this tool by fine-tuning BioBERT [12] using our in-house CDM (i.e., comprising 20 cohorts’ variable naming systems) in addition to two distinct CDMs. We defined a cross-connection among all three CDMs to drive a comprehensive variable embedding space from those. Finally, we enabled the AD-Mapper through a web interface to make the tool accessible.

## 2. Methods

### 2.1 Common data model

In developing a model that can distinguish between the semantics of different variables, two aspects played a major role. First, it was essential to include multiple ways that variables have been reported in different cohort studies and within the literature. For example, participants’ years of education were reported differently across cohort studies (Eduy, education, EDUC, etc.). Second, the inclusiveness of variable granularity was another vital aspect of establishing a successful model. The Apolipoprotein alleles (APOE) of participants were reported separately in certain datasets and together in other datasets. To address both of these factors, we consider a variety of ways that variables were addressed by including multiple cohorts’ naming systems. We utilized our in-house mapping CDM which consists of 20 distinct cohort datasets [2]. All cohorts’ variables were mapped to a reference term and an ontology where it was relevant. The reference terms were defined based on the variable description in addition to the abbreviation of the term where it was applicable (i.e., commonly used). For instance, the Mini-Mental State Examination is commonly referred to as MMSE, and as such, we defined the reference term as Mini-Mental State Examination (MMSE).

To expand the variability of the variable naming system, we manually harmonized previously constructed data models against our in-house model, namely, data models that were developed by Dementias Platform UK (DPUK) [7] and the Neuronet cohort initiative (NEURO Cohort). The set of harmonized variables generated by DPUK, called C-Surv, included 124 commonly measured variables among 4 distinct cohort studies. Similarly, the NEURO Cohort data model included 94 AD-related terms. Lastly, we harmonized our reference terms against the observational medical outcomes partnership (OMOP) CDM terms where the applicable term was available [14]. As a result, we created the AD-Mapper CDM.

### 2.2 The AD-Mapper NLP model

In this section, we explain the training, validation, and test data generated using the AD-Mapper CDM, the model training strategy of the AD-Mapper tool, and how the AD-Mapper can be used for inference. Additionally, we describe two different ways in which the mapping would be carried out: either using the reference terms (i.e., previously defined in AD-Mapper CDM) or using those reference terms in addition to prior knowledge of the cohorts’ and CDMs’ mappings.

#### 2.2.1 Training data

To investigate the feasibility of semantic harmonization of AD cohort studies using PLMs, we constructed a binary classification task aimed at discriminating between pairs of semantically equivalent variables (positives) and pairs that are not equivalent (negatives). By training on data consisting of positives and negatives, a fine-tuned PLM could generalize from it to find semantically equivalent pairs in a previously unseen set of variables. More specifically, we generated training data using the mappings that were established among different cohorts, CDMs, and the reference terms within the AD-Mapper CDM. We sampled different combinations of mappings among each reference term and mapped variables. For instance, the reference term ‘Age’ was mapped to multiple variables representing the age of participants in different cohorts and within CDMs (e.g., age, PTAGE, samplingAge, and age_at_visit, etc.). We created pairs for each mapping between ‘Age’ and all possible mapped variables. We followed a similar procedure for all pairs of mappings until we had generated one-to-one mappings, each labeled with 1 (i.e., positive) to represent semantically equivalent pairs.

For the negative mappings, we aimed to provide close variations of positive mapping pairs to enhance the model’s ability to distinguish very similar terms that are not equivalent. To achieve this, we created pairs of mappings within each modality, as variables grouped into a modality often had similar naming conventions. For instance, in the magnetic resonance imaging (MRI) mappings, we had left and right hippocampus volume and it was important for the model to learn the difference between ‘left’ and ‘right’. Therefore, we generated pairs that could be potentially confusing for the model to distinguish (e.g., Lhippo_FS_adj and Rhippo_FS_adj) and assigned them to class 0 (i.e., negative). This resulted in 13330 positive and 13330 negative labels. Moreover, we employed a weighted loss function where the negatives and positives are weighted by 0.1 and 0.9, respectively. This was undertaken to penalize false positives accurately and to represent the distribution of classes in later application scenarios, in which the majority of pairings between any two variables are expected to be non-semantically equivalent.

Variable mappings were frequently too uninformative for a model to learn the relation between the variables. To address this, we included descriptions where available for each variable in CDMs and cohort studies’ data dictionaries. For this purpose, we added two respective descriptions to each mapping pair in our training data. The description for the reference term was taken from the mapped ontology term. In contrast, the description for the mapped term was extracted from the corresponding data dictionary or the CDM from which the variable originated. Thereafter, we formed a sentence by concatenating each variable and its corresponding description (e.g., ‘variable’ + ‘its description’; **Figure 1, Step 2**). This strategy was then used for training our model. For further explanation, we refer to the Supplementary Material.

**Figure 1:**
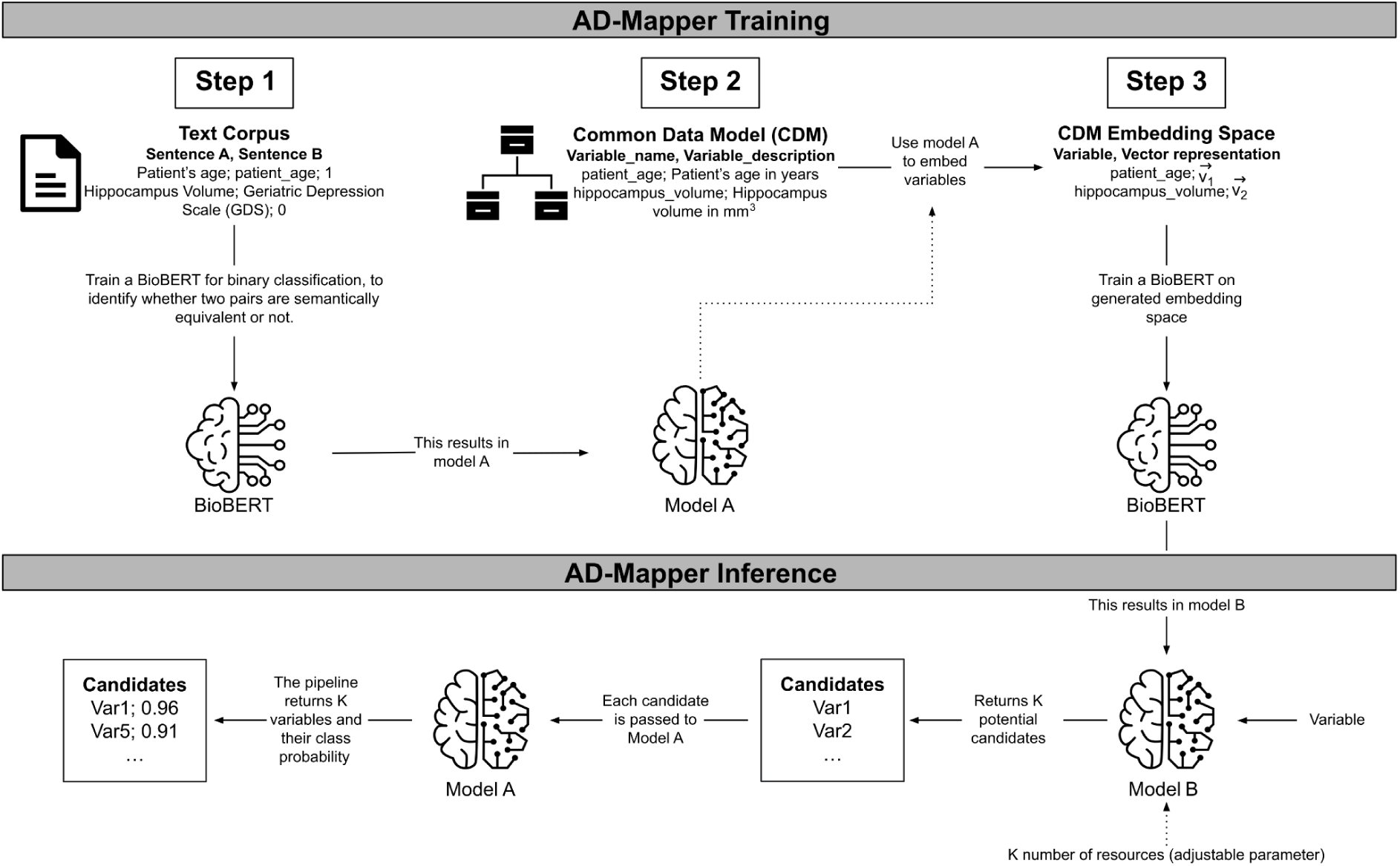
The underlying workflow of the AD-Mapper tool. Training the AD-Mapper consists of three steps: training BioBERT (i.e., Model A) on a text corpus, retrieving the names and descriptions from the CDM, and generating the embeddings. Then, the inference step comprises using Model B and A to generate a ranking of potential candidates and calculating the probabilities of positive mappings, respectively.

#### 2.2.2 Validation of cohort studies

For model tuning and performance investigation, we divided the dataset into train, test, and validation sets with fractions of 80/4/16, respectively. The test set consisted of 549 unseen examples. Additionally, to assess the performance of the plan strategy for the automatic harmonization of AD studies, we collected three distinct cohort studies (BRACE [15], AMED [16], and ALFA [17]) and manually harmonized them to the AD-Mapper CDM. The total number of available variables and mapping overlap between each cohort and the AD-Mapper CMD is shown in **Supplementary Table S1**. However, to evaluate the model’s performance on unseen datasets, we excluded these mappings from the training dataset.

#### 2.2.3 Models and training strategy

To develop a tool that could perform semantic harmonization of AD cohort studies, we developed a workflow that contained two models, Model A and B (**Figure 1, AD-Mapper Training**). First, we trained a Model A, performing a binary classification task using BioBERT [12] with a feed-forward classification head. This model compared two variables of the AD-Mapper CDM and their definitions, if available, and determined whether they were a match. Here, we considered two variables to be a match if they were semantically equivalent. In contrast, Model B, also based on BioBERT, was trained on embedding a variable into an embedding space (ℝ^768^) such that it was close, measured by the Euclidean distance, to its corresponding reference term. The initial embedding to be learned by model B was generated by taking each variable’s representation at the last hidden layer of BioBERT within model A. Initially, Model A and B were trained independently from each other, whereas they were used consecutively in the later application.

#### 2.2.4 Inference

The final pipeline employed in the AD-Mapper application consists of the two models (i.e., Models A and B) shown in **Figure 1** (i.e., AD-Mapper Inference). A mapping procedure starts with an input including a variable name and a suitable variable description if a description is available. This is concatenated to a joint sentence and fed into Model B. This first part of the pipeline returns K potential candidates, where K is a parameter provided by the user. Those candidates are the K closest variables in the embedding space where the variable gets mapped into while utilizing Model B. Then these K pairs, consisting of the input variable and each potential candidate, are fed into Model A, which returns a class probability for each pair indicating whether they belong to class 1, implying a match. All candidates below a certain threshold (e.g., 0.5) are eliminated and the rest is returned, where the one with the highest class probability is returned as the winner. The returned winner is the reference term suggested by the AD-Mapper.

As we previously established a mapping among all reference terms, cohort studies, and CDMs within the AD-Mapper CDM, we expanded the search for the best matches with this knowledge. We included another optional function, where not only the K candidates are determined by the model, but based on prior determined mappings, the model could also compare the new variable with those existing mappings. For instance, once K=5 candidates are generated as a result of model B, the total number of candidates fed to model A is enriched by all prior known mappings onto any of the 5 candidates. Finally, all 5 candidates and the prior known mappings onto those candidates are given to model A, and ultimately a winner is determined. In this case, the winner is either within one of the 5 original candidates or one of the later added ones. In the latter case, the final winner is determined by reversing the known mapping, and subsequently obtaining the reference term.

Considering that sometimes variables within cohort studies could potentially have a similar naming (e.g., AGE, age_at_visit), we added another optional functionality to the model to perform fuzzy string matching (https://github.com/maxbachmann/Levenshtein). We included this by assigning a weight to each methodology for the finalized mapping. This was done by introducing two weights *W*_1_, *W*_2_ ɛ [0, 1] where the first weight represents the BioBERT-based model (i.e., AD-Mapper model) and the second relies on the fuzzy string matching technique by calculating the Levenshtein distance between variables. This allows the user to decide whether the data should be harmonized using each technique separately or rather a weighted combination of both models.

### 2.3 The AD-Mapper Interface

The AD-Mapper application comes with two interfaces. The first one is a web-based graphical user interface (https://ad-mapper.scai.fraunhofer.de/), specifically designed for mapping single variables as well as .csv files. In both cases, the model requires variable names and their descriptions, if available, to perform the harmonization. Second, we provide powerful REST APIs that enable technically experienced users to employ the AD-Mapper in other applications. The APIs are fully documented and organized in a Swagger UI interface [18]. The interaction via REST APIs allows the user to fully leverage all configuration options that the mapping pipeline provides.

## 3. Results

### 3.1 AD-Mapper CDM

We constructed AD-Mapper CDM using previously established in-house data mappings and expanded the variables naming system by including three previously created CDMs. The AD-Mapper CDM included 20 cohorts’ variable naming systems semantically harmonized against a reference term and an ontology term per variable. In total 1218 unique reference terms were included in the AD-Mapper CDM. The total number of each cohort’s specific term and each external CDM that was harmonized against the reference terms is presented in **Table 1**. The overlap between our in-house CDM and the other external CDMs (i.e., C-Surv, NEURO Cohort, and OMOP) was small. One reason for this was that external CDMs were often developed based on the availability of measurements in this field, rather than the measurements themselves. To clarify further, certain terms were defined as biomarkers availability indicators (i.e., biomarker collected, yes or no) whereas AD-Mapper CDM focused on each specific measurement and how it was defined among different cohorts and CDMs. For instance, there were only 46 terms in C-Surv CDM that could be harmonized against our in-house CDM. Additionally, often CDMs consisted of a limited number of variables (e.g., NEURO Cohort 94 terms), which resulted in fewer terms being mapped as AD-Mapper CDM contained granular measurements. Lastly, we extended our AD-Mapper CDM to include previously unseen cohorts and additional variables. The total number of variables from each source is presented in **Table S2**.

**Table 1:** Total number of harmonized variables included in the AD-Mapper CDM. This table comprises all cohorts used for training the AD-Mapper model and hence excludes the ALFA, AMED, and BRACE cohorts. The final CDM presented on the AD-Mapper website consists of both training and test data as well as additional variables (resulting in 1300 unique variables, **Table S2**).

| Variable origin |  | Consortium | # Mapped variables |
| --- | --- | --- | --- |
| Cohort | **A4** [19] | Anti-Amyloid Treatment in Asymptomatic Alzheimer's Disease | 73 |
|  | **ABVIB** [20] | Aging Brain: Vasculature, Ischemia, and Behavior | 12 |
|  | **ADNI** [21] | The Alzheimer's Disease Neuroimaging Initiative | 340 |
|  | **AIBL** [22] | The Australian Imaging, Biomarker & Lifestyle Flagship Study of Ageing | 54 |
|  | **ANM** [23] | AddNeuroMed | 162 |
|  | **ARWIBO** [24] | Alzheimer's Disease Repository Without Borders | 1104 |
|  | **DOD-ADNI** [25] | Effects of TBI & PTSD on Alzheimer's Disease in Vietnam Vets | 322 |
|  | **EDSD** [26] | The European DTI Study on Dementia | 1061 |
|  | **EMIF** [27] | European Medical Information Framework | 30 |
|  | **EPAD** [28] | European Prevention of Alzheimer's Dementia | 116 |
|  | **I-ADNI** [29] | The Italian Alzheimer's Disease Neuroimaging Initiative | 1064 |
|  | **JADNI** [30] | Japanese Alzheimer's Disease Neuroimaging Initiative | 647 |
|  | **NACC** [31] | The National Alzheimer's Coordinating Center | 187 |
|  | **OASIS** [32] | Open Access Series of Imaging Studies | 1057 |
|  | **PREVENT-AD** [33] | Pre-symptomatic Evaluation of Experimental or Novel Treatments for Alzheimer's Disease | 34 |
|  | **PharmaCog** [34] | Prediction of Cognitive Properties of New Drug Candidates for Neurodegenerative Diseases in Early Clinical Development | 1067 |
|  | **ROSMAP** [35] | The Religious Orders Study and Memory and Aging Project | 29 |
|  | **VASCULAR** [36] | The Vascular Contributors to Prodromal Alzheimer's Disease | 53 |
|  | **VITA** [37] | Vienna Transdanube Aging | 1054 |
|  | **WMH-AD** [38] | White Matter Hyperintensities in Alzheimer's Disease | 1054 |
| **CDM** | **NEURO Cohort** | - | 14 |
|  | **C-Surv** [7] | - | 46 |
|  | **OMOP** [14] | The Observational Medical Outcomes Partnership | 107 |
| **Other** | **CURIE** [39] | Compact Uniform Resource Identifiers | 189 |
|  | **Reference term** | - | 1218 |

### 3.2 Model performance

We investigated the AD-Mapper’s performance by validating the model using the test split of the data, as well as distinct cohorts’ datasets that were excluded in the training step of the model. Using different weights and K values, we harmonized the test set and previously unseen datasets and calculated the accuracy of the mappings (**Table 2**). In addition, we compared the performance of the final pipeline with a baseline model. For that purpose, we used a string-matching model.

**Table 2:** Performance scores for harmonization of the test set and previously unseen cohorts using the AD-Mapper and string-matching as baseline comparison. K indicates the number of candidates to be assessed by the model.

| Dataset | Model | Prior knowledge | W-values<br>($W_1, W_2$) | K | Accuracy<br>(in %) |
| --- | --- | --- | --- | --- | --- |
| CDM test set | String-matching | - | - | - | 8.5 |
|  | AD-Mapper | - | 1.0, 0.0 | 1 | 72.9 |
|  | AD-Mapper | No | 1.0, 0.0 | 5 | 76.3 |
|  | AD-Mapper | Yes | 1.0, 0.0 | 5 | 74.88 |
|  | AD-Mapper | No | 1.0, 0.0 | 10 | 77.2 |
|  | AD-Mapper | Yes | 1.0, 0.0 | 10 | 73.93 |
| BRACE | String-matching | - | - | - | 26.4 |
|  | AD-Mapper | - | 1.0, 0.0 | 1 | 64.7 |
|  | AD-Mapper | No | 0.8, 0.2 | 5 | 76.4 |
|  | AD-Mapper | Yes | 0.8, 0.2 | 5 | 61.76 |
|  | AD-Mapper | No | 0.8, 0.2 | 10 | 76.4 |
|  | AD-Mapper | Yes | 0.8, 0.2 | 10 | 44.11 |
| AMED | String-matching | - | - | - | 4.16 |
|  | AD-Mapper | - | - | 1 | 59.09 |
|  | AD-Mapper | No | 1.0, 0.0 | 5 | 56.81 |
|  | AD-Mapper | Yes | 0.9, 0.1 | 5 | 70.45 |
|  | AD-Mapper | No | 1.0, 0.0 | 10 | 56.81 |
|  | AD-Mapper | Yes | 0.9, 0.1 | 10 | 61.36 |
| ALFA | String-matching | - | - | - | 15.51 |
|  | AD-Mapper | - | - | 1 | 67.92 |
|  | AD-Mapper | No | 0.7, 0.3 | 5 | 67.92 |
|  | AD-Mapper | Yes | 0.7, 0.3 | 5 | 54.71 |
|  | AD-Mapper | No | 0.7, 0.3 | 10 | 67.92 |
|  | AD-Mapper | Yes | 0.7, 0.3 | 10 | 54.71 |

Our results indicated that for the test dataset, the AD-Mapper had a superior accuracy of 77.2% while considering 10 candidates compared to the baseline approach (string-matching with 8.5% accuracy). Similarly, for all three cohort studies, previously unseen by the model, we observed that the AD-Mapper achieved (without incorporating prior known mappings) a much higher accuracy than the string-matching (BRACE 76.4% with K=5 and K=10; AMED 59.09% with K=1; and ALFA 67.92% with K=1, K=5 and K=10). Moreover, we noticed that, out of the cohorts harmonized, only the AMED cohort showed higher accuracy when using the model while utilizing prior known mappings than the one without. The AMED cohort had an accuracy of 70.45% while the model utilized the mappings of other cohort studies and CDMs with K=5. Furthermore, in all cases, except for the CDM test set, the model without prior knowledge achieved the same score for K=5 or K=10, which indicates that the model can predict the target variable from a small number of candidates.

We observed that different weights indicating the inclusion of different mapping strategies yielded various results. For instance, for the CDM test set, solely relying on the BioBERT-based prediction yielded the highest accuracy, while for the BRACE, AMED, and ALFA, combining the mapping technique (i.e., the BioBERT and string-matching) in a weighted manner reported better performance.

To further evaluate the AD-Mapper model, we exclusively utilized Model B and investigated whether the correct target variable is among the K candidates. The results of this analysis are presented in **Table S3**. In all instances where both K=5 and K=10 candidates were employed, they exhibited identical accuracy, except in the case of the test set. We observed the following accuracy rates: 71.7% for ALFA, 76.5% for BRACE, 79.5% for AMED, and 82.9% for the test set when using K=5, and 84.8% when using K=10. Additionally, we assessed whether it is possible to harmonize the variables that have not been included in the AD-Mapper CDM (**Table S4**). We provide additional details in the Supplementary Material.

### 3.3 Exemplary application scenario

We developed the AD-Mapper web interface and showcased our tool for accelerating data harmonization of AD cohort studies. Within this interface, users can simply upload their data dictionary as a .csv file and download the harmonized version. The interface allows users to select how the mapping should be carried out, either using the BioBERT-based model alone or in combination with the string-matching technique. An example of the AD-Mapper application scenario is illustrated in **Figure 2**. As shown in the figure, users can customize the weights and the number of candidates before executing the variable mapping. Users can also choose to have their data dictionary harmonized against the reference term or additionally have the harmonized version of all data sources available within the AD-Mapper CDM.

**Figure 2:**
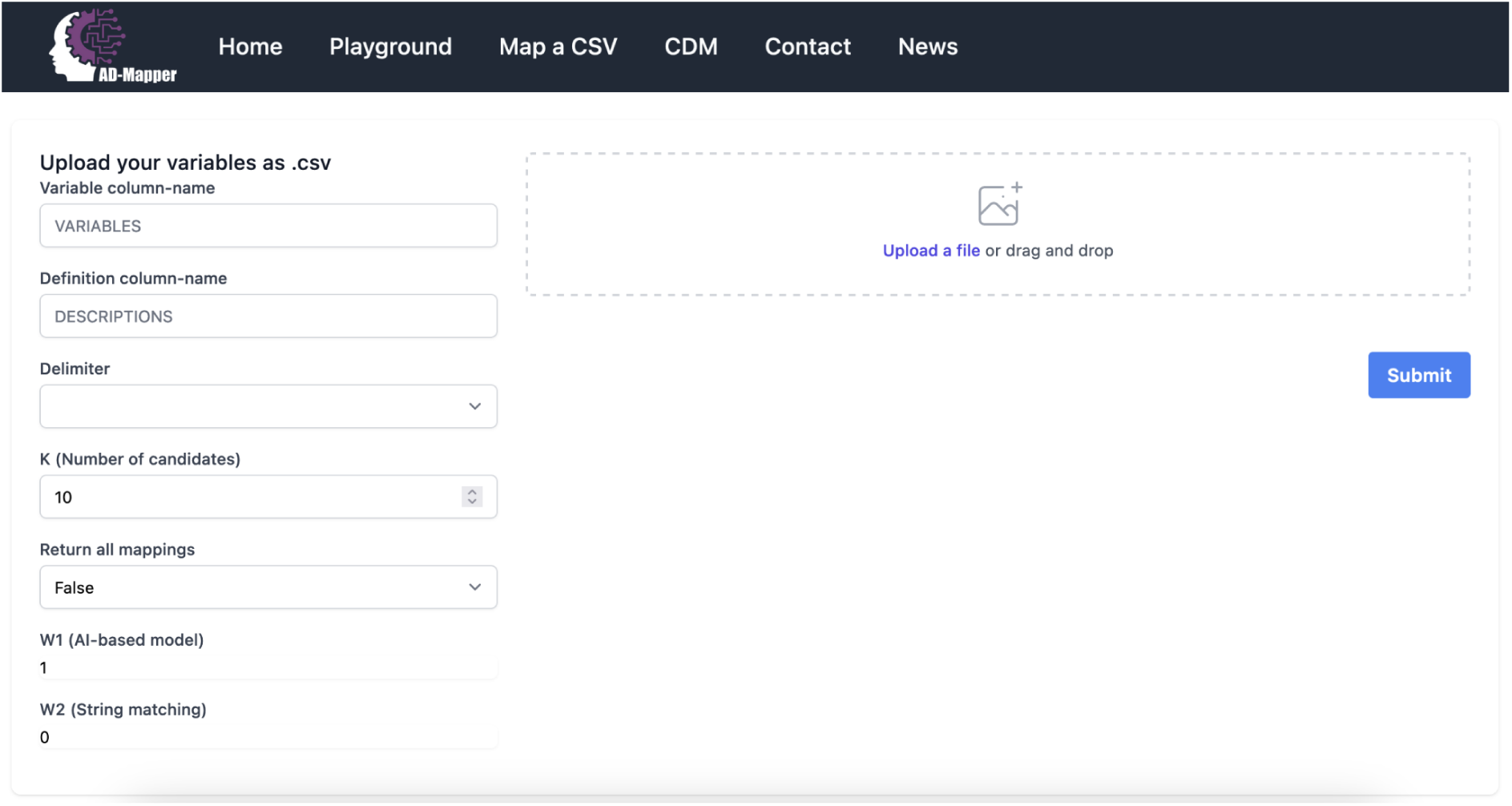
A page preview of the AD-Mapper user interface.

We included the AD-Mapper CDM as an additional function of the AD-Mapper interface. By doing so, we enable users to investigate cross-mappings among different cohorts and CDMs that have been harmonized against a reference term. Users can decide which weights are more suitable for their cohort dataset based on the similarity or dissimilarity of their data with the AD-Mapper CDM’s reference terms. For instance, when the variable naming system is defined similarly to our reference term, utilizing *W*_2_ (i.e., the string-matching technique) in addition to the BioBERT-based model could potentially result in higher accuracy. Lastly, users can easily download the AD-Mapper CDM to use it for data harmonization of the cohorts that were included and have been harmonized.

## 4. Discussion

In this work, we investigated whether semantic harmonization of cohort studies could be undertaken using automated models. Since data harmonization has frequently been a manual task, often very time-consuming, we explored the feasibility of employing a PLM to simplify this process. We fine-tuned a BioBERT model using a CDM that we generated, and evaluated the model’s performance using previously unseen datasets. Additionally, we compared our approach to a naive string-matching baseline model. Our results indicate that the AD-Mapper model can effectively facilitate the semantic harmonization of AD cohort studies.

### 4.1 Enabling variable transparency through a CDM

One underrepresented aspect of AD cohort studies is variable transparency and their naming conventions among cohorts or CDMs [2]. Even though multiple attempts were made previously to bring this aspect to light [8, 9, 2], most research focused on a limited number of variables. The AD-Mapper CDM addressed all of these challenges by considering a multitude of variables ranging from multiple modalities (1218 variables), and by including 23 different variable naming conventions (20 cohort studies and three CDMs). The AD-Mapper CDM can provide a valuable reference to highlight the underexplored biomarkers in this field. Another major concern was data privacy as data owners often prevent the researchers from uploading or sharing the data. We factored this aspect by using AD data dictionaries rather than the data itself as a foundation for our analysis and tools.

### 4.2 Automatic data harmonization

We evaluated the harmonization accuracy of unseen AD data using a naive approach as well as our proposed AD-Mapper model and showed that the latter technique exhibited superior accuracy. Furthermore, due to occasional similarities in naming systems among certain cohort studies, we investigated whether the inclusion of prior knowledge regarding cohort and CDM mappings would enhance the accuracy of correct match determination. This assessment demonstrated that, whilst the incorporation of prior knowledge increased harmonization accuracy for the AMED cohort, the opposite was observed in other cohorts. This could potentially be attributed to the AMED cohort sharing a highly similar naming system with the ADNI, DOD-ADNI, and JADNI cohort studies. By contrast, the other cohort studies did not share a similar naming system with any of the included variables’ mappings. Considering this finding, it can be inferred that employing prior knowledge could prove beneficial when similarities exist between the input cohort (i.e., the data requiring harmonization) and the variable naming system included within the AD-Mapper CDM, regardless of the cohort or CDM to which it closely corresponds.

To date, to our knowledge, there has been no automatic NLP-based harmonization of cohort studies conducted beyond standard string-matching techniques [10] or manual curation [40]. The application of such a standard technique (string-matching) led to relatively poor performance in the present study. This is highly important as utilizing cohort studies for data-driven investigation requires data harmonization and preprocessing [2, 6], and as such, incorrect harmonization of such cohorts could potentially lead to inaccurate discoveries. Although manual harmonization is often the most reliable approach, given the lengthy process of manual data harmonization, an automatic harmonization tool such as AD-Mapper could facilitate the procedure.

### 4.3 Model complexity and performance

To achieve better accuracy for the harmonization of cohort studies, we included a large number of variable naming systems stemming from different cohorts and CDMs within the AD-Mapper CDM. This factor influenced the variability of measurements being defined and subsequently resulted in 1218 unique reference terms. By selecting K candidates to explore and assess for finding the correct match for each unseen variable, the model expanded the search by that number of candidates to estimate the probability of them being a correct match. However, the choice of how many candidates are potentially sufficient to find the correct target for each unseen variable had a direct effect on the computational complexity of the model. Thus, the higher the number of candidates to be compared to, the higher the model complexity, affecting the inference speed. Despite this, our results revealed that the maximum accuracy was achieved by selecting K=5, and by increasing K to 10, the accuracy remained the same in the majority of cases. We observed the same results when solely utilizing Model B (see **Figure 1, Table S3**) to investigate whether the correct target could be found within a certain number of candidates. Taking this into account, we recommend that K=5 could be a sufficient standard for the number of candidates and that the model conducts the harmonization at a considerably quicker rate.

The accuracy of the variable harmonization was influenced by the choice of methodology that was utilized for finding the best mapping targets. One possible explanation is that by utilizing the string-matching technique in addition to the BioBERT-based mapping, the model is leveraging the similarities that exist between the potential candidates (the K-chosen reference terms) and the input variables to narrow down the best match. On the contrary, based on the observed result, we presume that the string-matching technique could decrease the accuracy of finding the best match when the input variables are not similar (i.e., they have a greater edit distance) to the reference terms. Thus, the ideal weights are highly dependent on the input cohort.

### 4.4 Limitations

One of the main limitations of our study was that we could only enable semantic harmonization, and the data distribution and measurement units may not be comparable across cohorts. This result stems from cohort studies employing certain exclusion and inclusion criteria while recruiting their participants, as well as differences in the way measurements were collected (e.g., different MRI devices). Additionally, cohort studies implement specific privacy agreements upon sharing the datasets, which hamper uploading or sharing of the data in any form (e.g., uploading data in the AD-Mapper interface). Given these challenges, within the scope of our paper, we could not achieve data interoperability beyond the semantics of variables. Here, we limited our mapping of the variables to those that are potentially comparable using a few preprocessing steps. Another limitation was that we could only cover the most commonly measured variables in our AD-Mapper CDM, and there are potentially certain variables that have not been included so far. This limitation was due to study-specific goals and the measurements collected to achieve these goals, as well as the granularity of variables shared with researchers. Future work can address this shortcoming by extending the AD-Mapper CDM, resulting in a refined embedding space, which ultimately leads to improved performance.

### 4.5 Conclusion

Data harmonization is an essential preliminary step in cross-cohort investigations before conducting data-driven analyses. Our objective was to expedite this often time-consuming process by developing the AD-Mapper interface. In doing so, we aimed to emphasize the importance of data compatibility and provide transparent insights into the biomarkers measured across diverse cohorts. The AD-Mapper demonstrated the feasibility of automatically harmonizing cohort studies, suggesting that this methodology could be applied to the study of different diseases.

## Code Availability

The code is available at: https://github.com/SCAI-BIO/ad-mapper

## Author Contributions

YS conceived and supervised the project. YS, MHA, and SB collected the datasets. YS and MCA harmonized the datasets. PW implemented the model and platform. YS and HB drafted the manuscript. LK, SB, and JG provided the CDMs. LK, SB, MCA, PW, JG, and MHA revised the manuscript. MHA acquired the funding. The authors read and approved the final manuscript.

## Abbreviations

A4: Anti-Amyloid Treatment in Asymptomatic Alzheimer’s Disease
ABVIB: Aging Brain Vasculature, Ischemia, and Behavior
AD: Alzheimer’s disease
ADNI: Alzheimer’s Disease Neuroimaging Initiative
AIBL: Australian Imaging, Biomarker & Lifestyle Flagship Study of Ageing
ALFA: For Alzheimer and Families
AMED: The Japanese Agency for Medical Research and Development
ANMerge: AddNeuroMed
APOE: Apolipoprotein Alleles
ARWIBO: Alzheimer’s Disease Repository Without Borders
BERT: Bidirectional Encoder Representations from Transformers
BioBERT: Bidirectional Encoder Representations from Transformers for Biomedical Text Mining
BRACE: Bristol Research into Alzheimer’s and Care for the Elderly
CDM: Common Data Model
DPUK: Dementias Platform UK
DOD-ADNI: Effects of TBI & PTSD on Alzheimer’s Disease in Vietnam Vets
DST: Data Stewardship Tool
EDSD: European DTI Study on Dementia
EMIF: European Medical Information Framework
EPAD: European Prevention of Alzheimer’s Dementia
I-ADNI: Italian Alzheimer’s Disease Neuroimaging Initiative
JADNI: Japanese Alzheimer’s Disease Neuroimaging Initiative
MMSE: Mini-Mental State Examination
MRI: Magnetic resonance imaging
NACC: National Alzheimer’s Coordinating Center
NLP: Natural Language Processing
OASIS: Open Access Series of Imaging Studies
OMOP: Observational Medical Outcomes Partnership
PREVENT-AD: Pre-symptomatic Evaluation of Experimental or Novel Treatments for Alzheimer’s Disease
PharmaCog: Prediction of Cognitive Properties of New Drug Candidates for Neurodegenerative Diseases in Early Clinical Development
PLM: Pre-trained Language Model (PLM)
ROSMAP: Religious Orders Study and Memory and Aging Project
VASCULAR: Vascular Contributors to Prodromal Alzheimer’s Disease
VITA: Vienna Transdanube Aging
WMH-AD: White Matter Hyperintensities in Alzheimer’s Disease

## Acknowledgments

We would like to express our appreciation to all data owners for their commitment to open science principles through the sharing of their data.

We thank the study participants and staff of the Rush Alzheimer’s Disease Center. ROSMAP was supported by NIA grants P30AG010161, R01AG015819, and R01AG017917.

The A4 Study is a secondary prevention trial in preclinical Alzheimer’s disease, aiming to slow cognitive decline associated with brain amyloid accumulation in clinically normal older individuals. The A4 Study is funded by a public-private-philanthropic partnership, including funding from the National Institutes of Health-National Institute on Aging, Eli Lilly and Company, Alzheimer’s Association, Accelerating Medicines Partnership, GHR Foundation, an anonymous foundation and additional private donors, with in-kind support from Avid and Cogstate. The companion observational Longitudinal Evaluation of Amyloid Risk and Neurodegeneration (LEARN) Study is funded by the Alzheimer’s Association and GHR Foundation. The A4 and LEARN Studies are led by Dr. Reisa Sperling at Brigham and Women’s Hospital, Harvard Medical School and Dr. Paul Aisen at the Alzheimer’s Therapeutic Research Institute (ATRI), University of Southern California. The A4 and LEARN Studies are coordinated by ATRI at the University of Southern California, and the data are made available through the Laboratory for Neuro Imaging at the University of Southern California. The participants screening for the A4 Study provided permission to share their de-identified data in order to advance the quest to find a successful treatment for Alzheimer’s disease. We would like to acknowledge the dedication of all the participants, the site personnel, and all of the partnership team members who continue to make the A4 and LEARN Studies possible. The complete A4 Study Team list is available on: a4study.org/a4-study-team.

Data collection and sharing of ABVIB was funded by the National Institutes on Aging (NIA) P01 AG12435.

Data collection and sharing of ARWIBO was supported by the Italian Ministry of Health, under the following grant agreements: Ricerca Corrente IRCCS Fatebenefratelli, Linea di Ricerca 2; Progetto Finalizzato Strategico 2000-2001 “Archivio normativo italiano di morfometria cerebrale con risonanza magnetica (età 40+)”; Progetto Finalizzato Strategico 2000-2001 “Decadimento cognitivo lieve non dementigeno: stadio preclinico di malattia di Alzheimer e demenza vascolare. Caratterizzazione clinica, strumentale, genetica e neurobiologica e sviluppo di criteri diagnostici utilizzabili nella realtà nazionale,”; Progetto Finalizzata 2002 “Sviluppo di indicatori di danno cerebrovascolare clinicamente significativo alla risonanza magnetica strutturale”; Progetto Fondazione CARIPLO 2005-2007 “Geni di suscettibilità per gli endofenotipi associati a malattie psichiatriche e dementigene”; “Fitness and Solidarietà”; and anonymous donors. Data used in the preparation of this article were obtained from the Alzheimer’s Disease Repository Without Borders (ARWiBo) (www.arwibo.it). The Principal Investigator of ARWIBO is Giovanni B.Frisoni, MD, University Hospitals and University of Geneva, Geneva, Switzerland, and IRCCS Fatebenefratelli, The National Centre for Alzheimer’s and Mental Diseases, Brescia, Italy. ARWIBO is the result of effort of many researchers of IRCCS Fatebenefratelli: G.Binetti, MD, Neurobiology; L.Bocchio-Chiavetto, PhD, Neuropharmacology; M.Cotelli, PhD, Neuropsychology Unit; C.Minussi, PhD, Neurophysiology; M.Gennarelli, PhD, Genetic Unit; R.Ghidoni, PhD, Proteomics Unit; D.Moretti, MD, and O.Zanetti, MD, Alzheimer’s Unit.

EPAD LCS is registered at www.clinicaltrials.gov Identifier: NCT02804789. Data used in preparation of this article were obtained from the EPAD LCS data set V.IMI, doi:10.34688/epadlcs_v.imi_20.10.30. The EPAD LCS was launched in 2015 as a public private partnership, led by Chief Investigator Professor Craig Ritchie MB BS. The primary research goal of the EPAD LCS is to provide a well-phenotyped probability-spectrum population for developing and continuously improving disease models for Alzheimer’s disease in individuals without dementia. This work used data and/or samples from the EPAD project which received support from the EU/EFPIA Innovative Medicines Initiative Joint Undertaking EPAD grant agreement n° 115736 and an Alzheimer’s Association Grant (SG21-818099-EPAD).

PharmaCog was funded through the European Community’s ‘Seventh Framework’ Programme (FP7/2007-2013) for an innovative scheme, the Innovative Medicines Initiative (IMI). IMI is a young and unique public-private partnership, founded in 2008 by the pharmaceutical industry (represented by the European Federation of Pharmaceutical Industries and Associations), EFPIA and the European Communities (represented by the European Commission).

J-ADNI was supported by the following grants: Translational Research Promotion Project from the New Energy and Industrial Technology Development Organization of Japan; Research on Dementia, Health Labor Sciences Research Grant; Life Science Database Integration Project of Japan Science and Technology Agency; Research Association of Biotechnology (contributed by Astellas Pharma Inc., Bristol-Myers Squibb, Daiichi-Sankyo, Eisai, Eli Lilly and Company, Merck-Banyu, Mitsubishi Tanabe Pharma, Pfizer Inc., Shionogi & Co., Ltd., Sumitomo Dainippon, and Takeda Pharmaceutical Company), Japan, and a grant from an anonymous Foundation.

Data collection and sharing for this project was funded by the Alzheimer’s Disease Neuroimaging Initiative (ADNI) (National Institutes of Health Grant U01 AG024904) and DOD ADNI (Department of Defense award number W81XWH-12-2-0012). ADNI is funded by the National Institute on Aging, the National Institute of Biomedical Imaging and Bioengineering, and through generous contributions from the following: AbbVie, Alzheimer’s Association; Alzheimer’s Drug Discovery Foundation; Araclon Biotech; BioClinica, Inc.; Biogen; Bristol-Myers Squibb Company; CereSpir, Inc.; Cogstate; Eisai Inc.; Elan Pharmaceuticals, Inc.; Eli Lilly and Company; EuroImmun; F. Hoffmann-La Roche Ltd and its affiliated company Genentech, Inc.; Fujirebio; GE Healthcare; IXICO Ltd.; Janssen Alzheimer Immunotherapy Research & Development, LLC.; Johnson & Johnson Pharmaceutical Research & Development LLC.; Lumosity; Lundbeck; Merck & Co., Inc.; Meso Scale Diagnostics, LLC.; NeuroRx Research; Neurotrack Technologies; Novartis Pharmaceuticals Corporation; Pfizer Inc.; Piramal Imaging; Servier; Takeda Pharmaceutical Company; and Transition Therapeutics. The Canadian Institutes of Health Research is providing funds to support ADNI clinical sites in Canada. Private sector contributions are facilitated by the Foundation for the National Institutes of Health (www.fnih.org). The grantee organization is the Northern California Institute for Research and Education, and the study is coordinated by the Alzheimer’s Therapeutic Research Institute at the University of Southern California. ADNI data are disseminated by the Laboratory for Neuro Imaging at the University of Southern California. This research was also supported by NIH grants P30 AG010129 and K01 AG030514.

The NACC database is funded by NIA/NIH Grant U24 AG072122. NACC data are contributed by the NIA-funded ADRCs: P30 AG062429 (PI James Brewer, MD, PhD), P30 AG066468 (PI Oscar Lopez, MD), P30 AG062421 (PI Bradley Hyman, MD, PhD), P30 AG066509 (PI Thomas Grabowski, MD), P30 AG066514 (PI Mary Sano, PhD), P30 AG066530 (PI Helena Chui, MD), P30 AG066507 (PI Marilyn Albert, PhD), P30 AG066444 (PI John Morris, MD), P30 AG066518 (PI Jeffrey Kaye, MD), P30 AG066512 (PI Thomas Wisniewski, MD), P30 AG066462 (PI Scott Small, MD), P30 AG072979 (PI David Wolk, MD), P30 AG072972 (PI Charles DeCarli, MD), P30 AG072976 (PI Andrew Saykin, PsyD), P30 AG072975 (PI David Bennett, MD), P30 AG072978 (PI Neil Kowall, MD), P30 AG072977 (PI Robert Vassar, PhD), P30 AG066519 (PI Frank LaFerla, PhD), P30 AG062677 (PI Ronald Petersen, MD, PhD), P30 AG079280 (PI Eric Reiman, MD), P30 AG062422 (PI Gil Rabinovici, MD), P30 AG066511 (PI Allan Levey, MD, PhD), P30 AG072946 (PI Linda Van Eldik, PhD), P30 AG062715 (PI Sanjay Asthana, MD, FRCP), P30 AG072973 (PI Russell Swerdlow, MD), P30 AG066506 (PI Todd Golde, MD, PhD), P30 AG066508 (PI Stephen Strittmatter, MD, PhD), P30 AG066515 (PI Victor Henderson, MD, MS), P30 AG072947 (PI Suzanne Craft, PhD), P30 AG072931 (PI Henry Paulson, MD, PhD), P30 AG066546 (PI Sudha Seshadri, MD), P20 AG068024 (PI Erik Roberson, MD, PhD), P20 AG068053 (PI Justin Miller, PhD), P20 AG068077 (PI Gary Rosenberg, MD), P20 AG068082 (PI Angela Jefferson, PhD), P30 AG072958 (PI Heather Whitson, MD), P30 AG072959 (PI James Leverenz, MD).

The results published here are in whole or in part based on data obtained from the AD Knowledge Portal (https://adknowledgeportal.org). These data are from the Vascular Contributors to Prodromal Alzheimer’s disease (VASCULAR) study. The study is part of the M2OVE-AD supported studies and is supported by NIH/NIA (PI: Ihab Hajjar, MD, MS grant numbers: AG051633 and AG057470. The investigators acknowledge the contribution of the study participants who donated their time and effort for this study.

