## Supplementary Material for "Semantic harmonization of Alzheimer’s disease datasets using AD-Mapper"

* Data used in preparation of this article were obtained from the Alzheimer's Disease Neuroimaging Initiative (ADNI) database (adni.loni.usc.edu). As such, the investigators within the ADNI contributed to the design and implementation of ADNI and/or provided data but did not participate in analysis or writing of this report. A complete listing of ADNI investigators can be found at: http://adni.loni.usc.edu/wp-content/uploads/how_to_apply/ADNI_Acknowledgement_List.pdf

† Japanese Alzheimer's Disease Neuroimaging Initiative: Data used in preparation of this article were obtained from the Japanese Alzheimer's Disease Neuroimaging Initiative (J-ADNI) database deposited in the National Bioscience Database Center Human Database, Japan (Research ID: hum0043.v1, 2016). As such, the investigators within J-ADNI contributed to the design and implementation of J-ADNI and/or provided data but did not participate in analysis or writing of this report. A complete listing of J-ADNI investigators can be found at: https://humandbs.biosciencedbc.jp/ en/hum0043-j-adni-authors.

‡ Data used in preparation of this article were obtained from the Aging Brain: Vasculature, Ischemia, and Behavior Study (ABVIB). As such, the key investigators within the ABVIB contributed to the design and implementation of ABVIB and/or provided data but did not participate in analysis or writing of this report: Helena C. Chui M.D. (Principal Investigator), Charles C. DeCarli, M.D., William G. Ellis, M.D., William J. Jagust, M.D., Joel H. Kramer, Ph.D., Meng Law, M.D., Dan Mungas Ph,D. Bruce R. Reed, Ph.D., Nerses Sanossian, M.D., Michael W. Weiner, M.D. Wendy J. Mack, Ph.D., Harry V. Vinters, M.D., Chris Zarow, Ph.D., Ling Zheng, Ph.D.

§ Data used in preparation of this article were obtained from the Alzheimer's Disease Repository Without Borders (ARWiBo) database (www.arwibo.it). As such, the researchers within the ARWiBo contributed to the design and implementation of ARWiBo and/or provided data but did not participate in analysis or writing of this report. A complete listing of ARWiBo researchers can be found at: www.arwibo.it/acknowledgement.it

‡ Data used in preparation of this article were obtained from the Longitudinal Cohort Study (LCS), delivered by the European Prevention of Alzheimer's Disease (EPAD) Consortium. As such investigators within the EPAD LCS and EPAD Consortium contributed to the design and implementation of EPAD and/or provided data but did not participate in analysis or writing of this report. A complete list of EPAD Investigators can be found at: http://ep-ad.org/wp-content/uploads/2020/12/202010_List-of-epadistas.pdf

| Dataset | Consortium | # Overlapping variables | # Total variables |
|---|---|---|---|
| BRACE [1] | Bristol Research into Alzheimer's and Care for the Elderly | 34 | 476 |
| AMED [2] | The Japanese Agency for Medical Research and Development | 44 | 691 |
| ALFA [3] | For Alzheimer and Families | 53 | 252 |

**Table S1:** The total number of collected variables in BRACE, AMED, and ALFA studies as well as the number of variables (i.e., the number of variables that could be harmonized against the AD-Mapper).

| Variable origin | | # Mapped variables |
|---|---|---|
| **Cohort** | **A4** [4] | 97 |
| | **ABVIB** [5] | 12 |
| | **ADNI** [6] | 413 |
| | **AIBL** [7] | 58 |
| | **ALFA** [3] | 56 |
| | **AMED** [2] | 105 |
| | **ANM** [8] | 162 |
| | **ARWIBO** [9] | 1129 |
| | **BRACE** [1] | 36 |
| | **DOD-ADNI** [10] | 395 |
| | **EDSD** [11] | 1061 |
| | **EMIF** [12] | 31 |
| | **EPAD** [13] | 140 |
| | **I-ADNI** [14] | 1070 |
| | **JADNI** [15] | 720 |
| | **NACC** [16] | 229 |
| | **OASIS** [17] | 1057 |
| | **PREVENT-AD** [18] | 35 |

| | | |
|---|---|---|
| | **PharmaCog** [19] | 1073 |
| | **ROSMAP** [20] | 30 |
| | **VASCULAR** [21] | 55 |
| | **VITA** [22] | 1054 |
| | **WMH-AD** [23] | 1054 |
| **CDM** | **NEURO Cohort** | 14 |
| | **C-Surv** [24] | 47 |
| | **OMOP** [25] | 144 |
| **Other** | **CURIE** [26] | 218 |
| | **Reference term** | 1300 |

**Table S2:** Total number of mapped variables in the extended AD-Mapper CDM.

| Dataset | K | Accuracy (in %) |
|---|---|---|
| CDM test set | 5 | 82.93 |
| | 10 | 84.83 |
| BRACE | 5 | 76.47 |
| | 10 | 76.47 |
| AMED | 5 | 79.54 |
| | 10 | 79.54 |
| ALFA | 5 | 71.69 |
| | 10 | 71.69 |

**Table S3:** Performance scores of utilizing the Model B alone. K indicates the number of candidates to be assessed by the model.

### Variables that fall outside the AD-Mapper common data model (CDM)

Since cohort studies were conducted to address specific research questions, the measurements collected often vary from one cohort to another. In the AD-Mapper CDM, our goal was to include variables that were common in at least two cohorts. Similarly, as mentioned in Section 3.1 of the main manuscript, the total number of included variables in the external CDMs was limited. Consequently, the variable naming space of our CDM was limited to those cohorts and CDMs. Given this challenge, a highly relevant question was how the model would handle variables that were not initially part of the AD-Mapper CDM and had not yet been incorporated into the learned embedding space. To assess the pipeline's performance in this scenario, we manually mapped 82 variables from the ADNI cohort and added them to the AD-Mapper CDM, after which they were integrated into the embedding space. Finally, we conducted an experiment to evaluate the model's accuracy in mapping variables that were added to the model at a later stage. For this analysis, we used Model A (classifier) to estimate the accuracy of harmonizing the Alzheimer's Disease Neuroimaging Initiative (ADNI) cohort. Subsequently, we performed a similar analysis using the string-matching technique (**Table S3**). Our results revealed an accuracy of 48.1% compared to the string-matching accuracy of 12.1%.

| **Model** | **W-values ($W_1$, $W_2$)** | **K** | **Accuracy (in %)** |
|---|---|---|---|
| String-matching | - | - | 12.1 |
| AD-Mapper | 1.0,0.0 | 82 | 48.05 |

**Table S4:** Performance on completely unseen variables of ADNI harmonized using the AD-mapper and string-matching techniques.

#### Importance of data dictionary descriptiveness

As described in Section 2.2.1 of the manuscript, we utilized the mappings of different cohorts and common data models (CDMs) to generate a training dataset. We included variable descriptions wherever available to enhance the model's comprehension of the variables' semantics. This step was taken because variables in cohort studies frequently lacked descriptiveness, and by adding these descriptions, the model could potentially identify similarities between the input variables and the reference terms. Similarly, when the model was used for harmonization of the new cohort studies (i.e., previously unseen by the model and thus, excluded from the AD-Mapper CDM), the variable descriptions played a significant role. However, in the version of the AMED cohort's data dictionary that we retrieved, the variable descriptions were not readable and displayed question marks, possibly due to a formatting issue. Considering the similarities between the variable naming convention of AMED and those of ADNI, JADNI, I-ADNI, and

DOD-ADNI cohorts, the model could potentially have performed better if the descriptions for all variables in the AMED cohort were available.
